# Lipid composition effects on the number and size of liposomes formed by the inverted emulsion method

**DOI:** 10.1101/2025.07.07.663091

**Authors:** Hibiki Sakata, Hitomi Matubara, Kanako Gomi, Makito Miyazaki

## Abstract

Liposomes have been widely employed as membrane scaffolds in the construction of minimal cell models. In 2002, the inverted emulsion method was introduced as a novel technique for generating giant liposomes by transferring water-in-oil droplets across an oil/water interface. This technique enables the encapsulation of purified proteins or cytoplasmic extracts into cell-sized liposomes under physiological buffer conditions, and has since become a cornerstone in bottom-up synthetic biology. Despite its broad application over the past two decades, the effects of lipid composition on the production yield and size distribution of liposomes generated by the inverted emulsion method remain largely unknown. In this study, we systematically investigated the effects of phospholipid composition on the production yield and size distribution of liposomes generated using the inverted emulsion method. We used a natural phosphatidylcholine purified from chicken egg yolk (egg PC) as the base membrane component, and examined the extent to which substituting a fraction of egg PC with other phospholipids—differing in net electric charge and/or the molecular weight of their hydrophilic head groups—affects the number and size distribution of liposomes. We found that a 10% replacement of egg PC with charged phospholipids significantly enhances the production yield of liposomes by approximately tenfold, transformed by both the natural gravitational force and the greater force applied by centrifugation. In addition, lipids with smaller head groups slightly enhance the production yield in both methods. Lipids with larger head groups also slightly enhance the production yield and promote the formation of larger liposomes in the natural sedimentation method, but these effects are diminished in the centrifugal sedimentation method, presumably due to the strong centrifugal force. These findings provide valuable guidelines for optimizing preparation protocols for minimal cell models.

## INTRODUCTION

How the cell, the smallest unit of life, emerges from molecules is a central question in biology. To tackle this fundamental yet extremely challenging question, scientists have put great effort into creating minimal artificial cell models by encapsulating molecules (proteins and nucleic acids) into liposomes and reconstituting biological functions from a minimal set of components <u>(Miyata 1992, Takiguchi 2008, Yanagisawa 2011, Tsai 2011, Takiguchi 2011, Carvalho 2013, Peters 2015, Loiseau 2016, Nies 2018, Dürre 2018, Berhanu 2019, Buddingh 2020, Bashirzadeh 2021, Razavi 2024, Sakamoto 2024, Monck 2024, Matsubayashi 2024, Sakamoto 2024, Takamori 2025)</u>. The early method for preparing liposomes is the hydration method, in which dried lipid films are hydrated by an aqueous solution, and the hydrated lipid layers spontaneously assemble into liposomes during the peeling process <u>(Reeves 1969, Walde 2010, Morales-Penningston 2010)</u>. However, the efficiency of vesicle formation is highly sensitive to the composition of the hydration solution. In particular, encapsulating proteins under physiological buffer conditions is challenging <u>(Tsai 2011)</u>, which is a critical limitation for constructing artificial cell models. Moreover, the lamellarity of these liposomes varies, leading to contamination by multilamellar vesicles <u>(Akashi 1996, Tsai 2011, Chiba 2014)</u>. This limitation was addressed to some extent with the development of the electroformation method in the 1980s, which uses an electric field to produce more uniform liposomes <u>(Angelova 1986)</u>. However, this method also has limitations regarding the buffer conditions required for efficient liposome formation. Furthermore, the low encapsulation efficiency of proteins has hindered progress in synthetic biology research.

In 2003, Pautot *et al*. developed a novel method to produce liposomes by transferring an inverted emulsion, *i.e.*, water-in-oil droplets, across the water/oil interface, assisted by the density difference between oil and water <u>(Pautot 2003)</u> (**Fig. 1A**). The authors named this technique the inverted emulsion method, which is also referred to by some researchers as the droplet transfer method. This method significantly improved the efficiency of encapsulating proteins into cell-sized liposomes under physiological buffer conditions. In addition, this method enables the production of unilamellar liposomes with high probability, exhibiting strong robustness to variations in lipid composition and concentration <u>(Chiba 2014)</u>. Owing to these advantages, this method has been extensively used to encapsulate proteins of interest or cell extracts, including cell-free protein expression systems, into liposomes to create artificial cell models <u>(Noireaux 2004, Maeda 2012, Berhanu 2019)</u>. In the original method, centrifugation was used to transfer the droplets into liposomes to enhance the transfer and sedimentation rates. Many biophysical studies and synthetic biology studies followed this protocol to encapsulate proteins or cell extracts <u>(Noireaux 2004, Takiguchi 2008, Yanagisawa 2011, Tsai 2011, Takiguchi 2011, Maeda 2012, Carvalho 2013, Peters 2015, Loiseau 2016, Nies 2018, Dürre 2018, Berhanu 2019, Buddingh 2020, Bashirzadeh 2021, Razavi 2024, Sakamoto 2024, Monck 2024, Matsubayashi 2024, Sakamoto 2024, Takamori 2025)</u>. Alternatively, several studies have used gravitational force to spontaneously transfer the droplets into liposomes <u>(Takiguchi 2008, Yanagisawa 2011, Takiguchi 2011, Ito 2013)</u>. This natural sedimentation method offers a simpler and more straightforward manipulation protocol compared to the centrifugal sedimentation method, enabling the observation of internal reactions and membrane deformation processes immediately after the transformation of droplets into liposomes under an optical microscope <u>(Takiguchi 2008, Takiguchi 2011)</u>.

**Figure 1.**
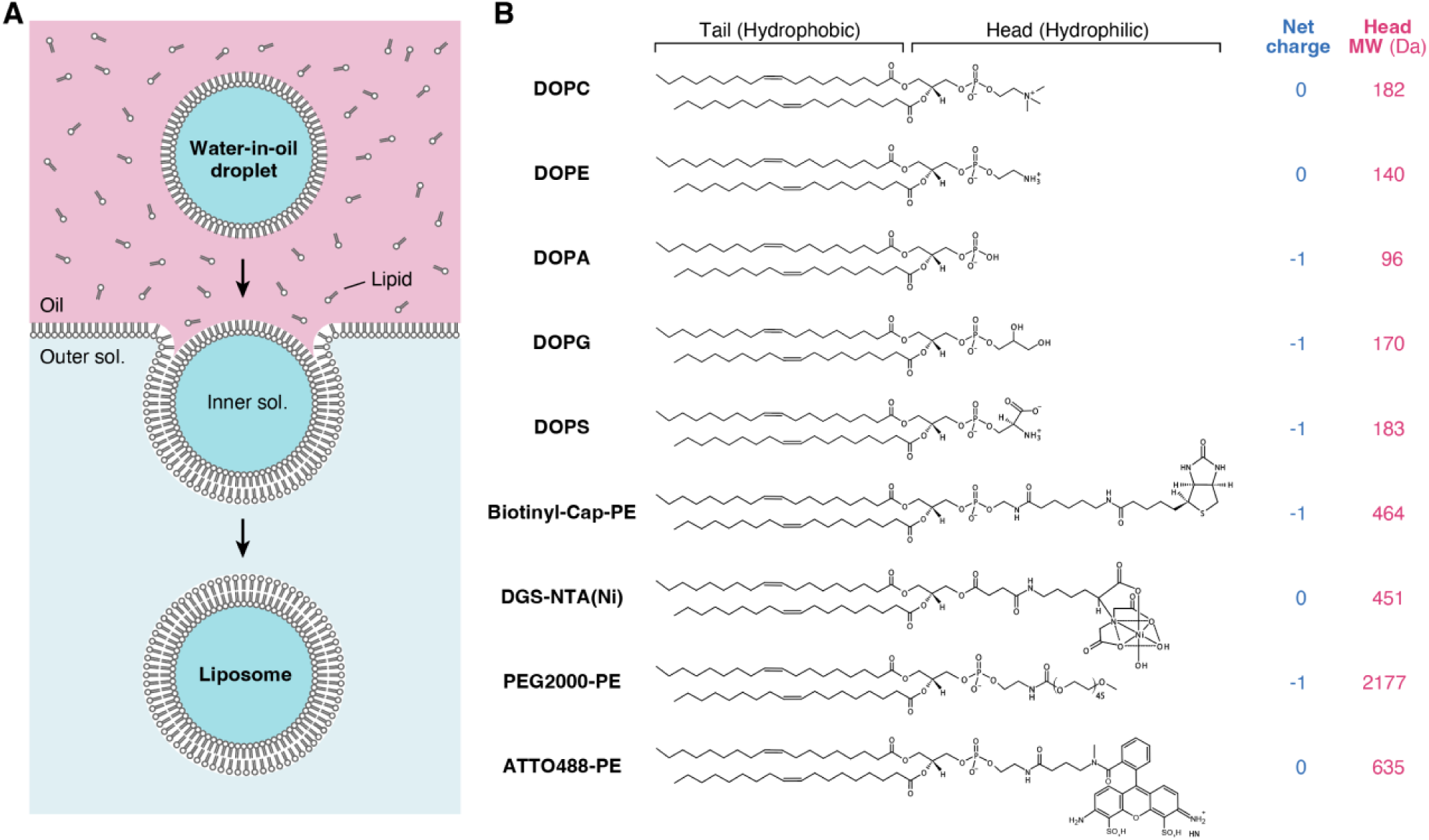
Liposome formation method and chemical structures of phospholipids. **A.** Schematic illustration of the inverted emulsion method for liposome formation. Water-in-oil droplets surrounded by a lipid monolayer were transformed into giant liposomes through natural sedimentation (**Figs. 2 and 3**) or centrifugation (**Figs. 4, 5, and 6**). Hereafter, these methods will be referred to as the “natural sedimentation method” and the “centrifugal sedimentation method,” respectively. **B.** Chemical structures of the phospholipids used in this study. The net electric charges and molecular weights of the hydrophilic regions are listed on the left.

More than two decades have passed since the invention of the inverted emulsion technique. Despite its extensive use in various biophysical and synthetic biology studies, preparing protein-encapsulated liposomes with high efficiency remains challenging. Specifically, it is generally agreed among researchers that the number of liposomes produced by the inverted emulsion method is small, limiting experimental efficiency. This low production yield becomes a critical issue when attempting to encapsulate highly concentrated proteins or membrane-associated proteins. To overcome this problem, several derivative methods, including the continuous droplet interface crossing encapsulation (cDICE) method <u>(Abkarian 2011)</u> and a cDICE-based simpler method <u>(Bashirzadeh 2021)</u>, have been proposed, and utilized for synthetic biology studies <u>(Loiseau 2016, Dürre 2018)</u>. However, these methods require a specially designed device for liposome production. As a simpler and more accessible strategy, researchers have empirically added negatively charged lipids, such as phosphatidylglycerol (PG), to phosphatidylcholine (PC), a neutrally charged lipid used as the base membrane component. Nevertheless, to the best of our knowledge, a systematic quantitative study examining the effects of lipid composition on the number and size of liposomes produced by the inverted emulsion method is still lacking.

In this study, we systematically investigate the effects of phospholipid composition (**Fig. 1B**) on the production yield and size distribution of liposomes formed by both natural sedimentation and centrifugal sedimentation methods. We use egg PC, a natural PC purified from chicken egg yolk, as the base membrane component, and examine to what extent replacing a fraction of egg PC with other phospholipids exhibiting different chemical properties (net electric charge) and/or having a different molecular weight of the hydrophilic part affects the number and size distribution of liposomes.

## MATERIALS AND METHODS

### Lipids

L-α-phosphatidylcholine from chicken egg yolk (egg PC; 840051P), 1,2-dioleoyl-sn-glycero-3-phosphocholine (DOPC; 850375P), 1,2-dioleoyl-sn-glycero-3-phosphoethanolamine (DOPE; 850725P), 1,2-dioleoyl-sn-glycero-3-phosphate (sodium salt) (DOPA; 840875P), 1,2-dioleoyl-sn-glycero-3-phosphatidylglycerol (DOPG; 840475P), 1,2-dioleoyl-sn-glycero-3-phospho-L-serine (sodium salt) (DOPS; 840035P), 1,2-dioleoyl-sn-glycero-3-phosphoethanolamine-N-(cap biotinyl) (sodium salt) (Biotinyl-Cap-PE, 870273), 1,2-dioleoyl-sn-glycero-3-[(N-(5-amino-1-carboxypentyl)iminodiacetic acid)succinyl] (nickel salt) (DGS-NTA(Ni); 790404C), and 1,2-dioleoyl-sn-glycero-3-phosphoethanolamine-N-[methoxy(polyethylene glycol)-2000] (ammonium salt) (PEG2000-PE: 880130P) were purchased from Avanti Polar Lipids. 1,2-Dioleoyl-*sn*-glycero-3-phosphoethanolamine labeled with Atto 488 (ATTO488-PE; 488-16) was purchased from ATTO-TEC. All lipids were used without further purification. Since egg PC is a mixture of phosphatidylcholines with various unsaturated and saturated fatty acids, the mean molecular weight (770.123 Da) was used to calculate the molar concentration (**Fig. S1**).

### Buffers

All experiments were performed using A50 buffer (50 mM HEPES-KOH pH 7.6, 50 mM KCl, 5 mM MgCl_2_, 1 mM EGTA) containing 150 mM sucrose and 350 mM glucose for the inner solution, and A50 buffer containing 500 mM glucose for the outer solution, respectively. To visualize liposomes under an epi-fluorescence microscope, 10 μM of octadecyl rhodamine B chloride (R18) (O246, Invitrogen) was added to the inner solution as needed.

### Preparation of lipid-oil mixture

The lipid-oil mixture was prepared according to our previous report <u>(Chiba 2014)</u>. Briefly, lipids were dissolved in chloroform, which was dehydrated with molecular sieves 4A, at a concentration of 20 mM. The lipid solutions were mixed at a desired molar ratio and the mixture was put into a 1.5 mL glass test tube (0407-03, Maruemu). Then, the tube was placed in a vacuum desiccator overnight to evaporate chloroform. Next day, the lipid dry film formed at the bottom of the test tube was dissolved in 1 mL of mineral oil (M8410, Sigma-Aldrich for **Figs. 1,2, 3, and 6** or 23306-84, Nacalai Tesque for **Figs. 4 and 5**) using a heat bath at 80°C for 30 min, followed by using a bath sonicator (AS12GTU, As One) at 60°C and a power of 60 W for 90 min. The lipid-oil mixture was stored at room temperature under dark conditions overnight and used the next day. The total lipid concentration in oil was fixed at 1 mM for all the experiments.

**Figure 2.**
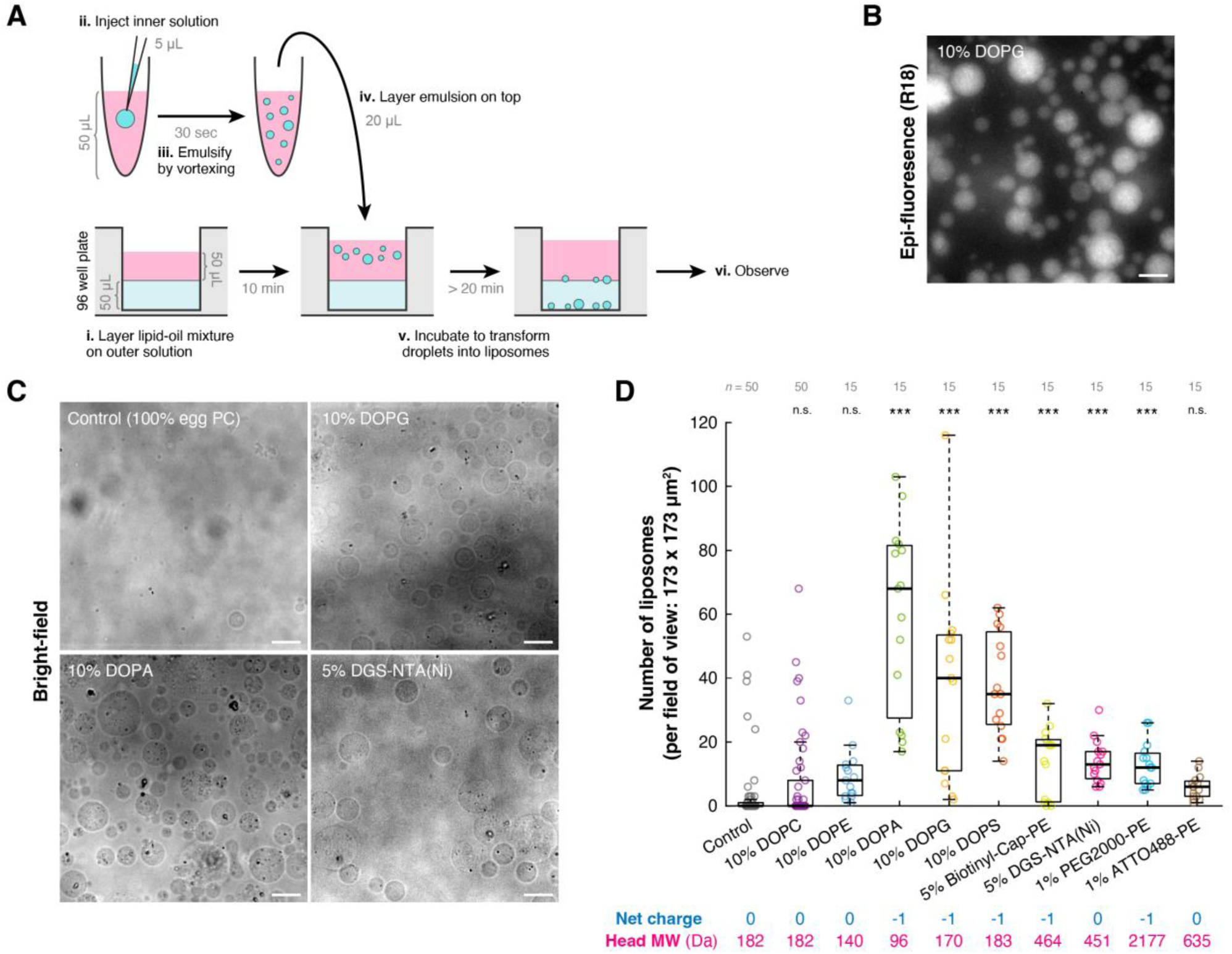
Effects of lipid composition on the production yield of liposomes formed using the natural sedimentation method. **A.** Schematic illustration of the natural sedimentation method for liposome formation and observation. **B.** Representative epi-fluorescence image of liposomes. The corresponding bright-field image of the same field of view is shown in **C,** top-right panel. Scale bar: 20 μm. **C.** Representative bright-field images of liposomes. Scale bars: 20 μm. **D.** Number of liposomes observed in a field of view. The raw data are overlaid on the box plot as a scatter plot. In each experiment, five fields of view were observed, and the number of liposomes larger than 5 μm in diameter was counted in each field. Three to ten independent experiments were performed for each condition. The net electric charges and molecular weights of the hydrophilic regions of the additive phospholipids or egg PC (control) are listed. ***: *p* < 0.001, n.s.: *p* ≥ 0.05 (Welch’s *t*-test, two-sided).

**Figure 3.**
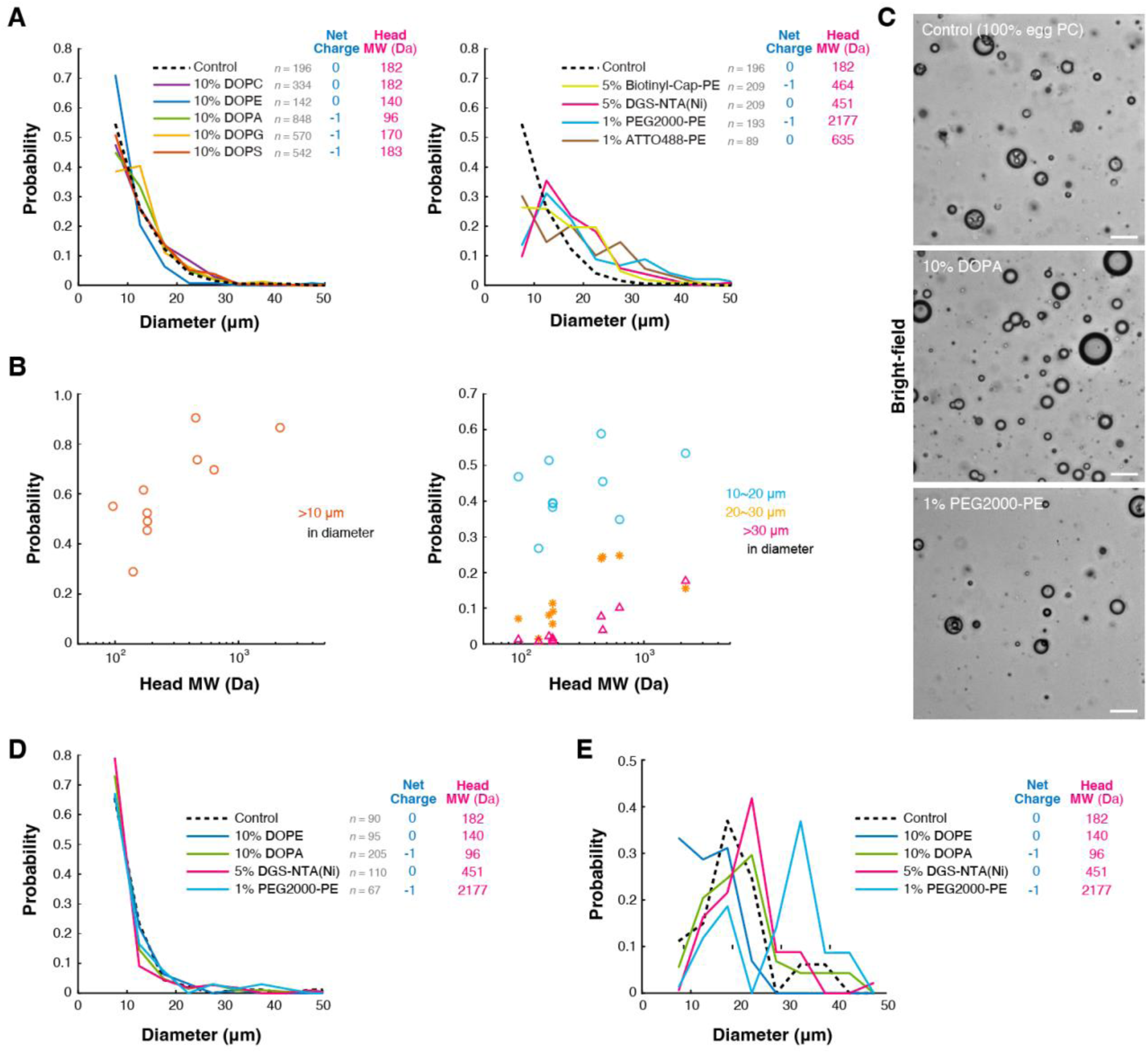
Effects of lipid composition on the diameter of liposomes formed using the natural sedimentation method. **A.** Distribution of liposome diameters. Three to ten independent experiments were performed for each condition. Left: Smaller head additive lipids. Right: Larger head additive lipids. The net electric charges and molecular weights of the hydrophilic regions of the additive phospholipids or egg PC (control) are listed. **B.** Relationship between the molecular weight of the hydrophilic regions of the additive phospholipids and liposome diameter. Left: Probabilities of liposomes larger than 10 μm in diameter (orange circles). Right: Probabilities of liposomes between 10 μm and 20 μm in diameter (blue circles), liposomes between 20 μm and 30 μm in diameter (yellow asterisks), and liposomes larger than 30 μm in diameter (magenta triangles) are compared. **C.** Representative images of water-in-oil droplets observed using a bright-field microscope. Scale bars: 20 μm. **D.** Distribution of water-in-oil droplet diameters. Two independent experiments were performed for each condition. The net electric charges and molecular weights of the hydrophilic regions of the additive phospholipids or egg PC (control) are listed. **E.** Droplet-to-liposome transformation efficiency. The probability distribution of liposome diameter (**Fig. 3A**) was divided by that of droplet diameter (**Fig. 3D**) and normalized against the diameter. A higher efficiency indicates a higher transformation rate of water-in-oil droplets into liposomes at the oil/water interface.

**Figure 4.**
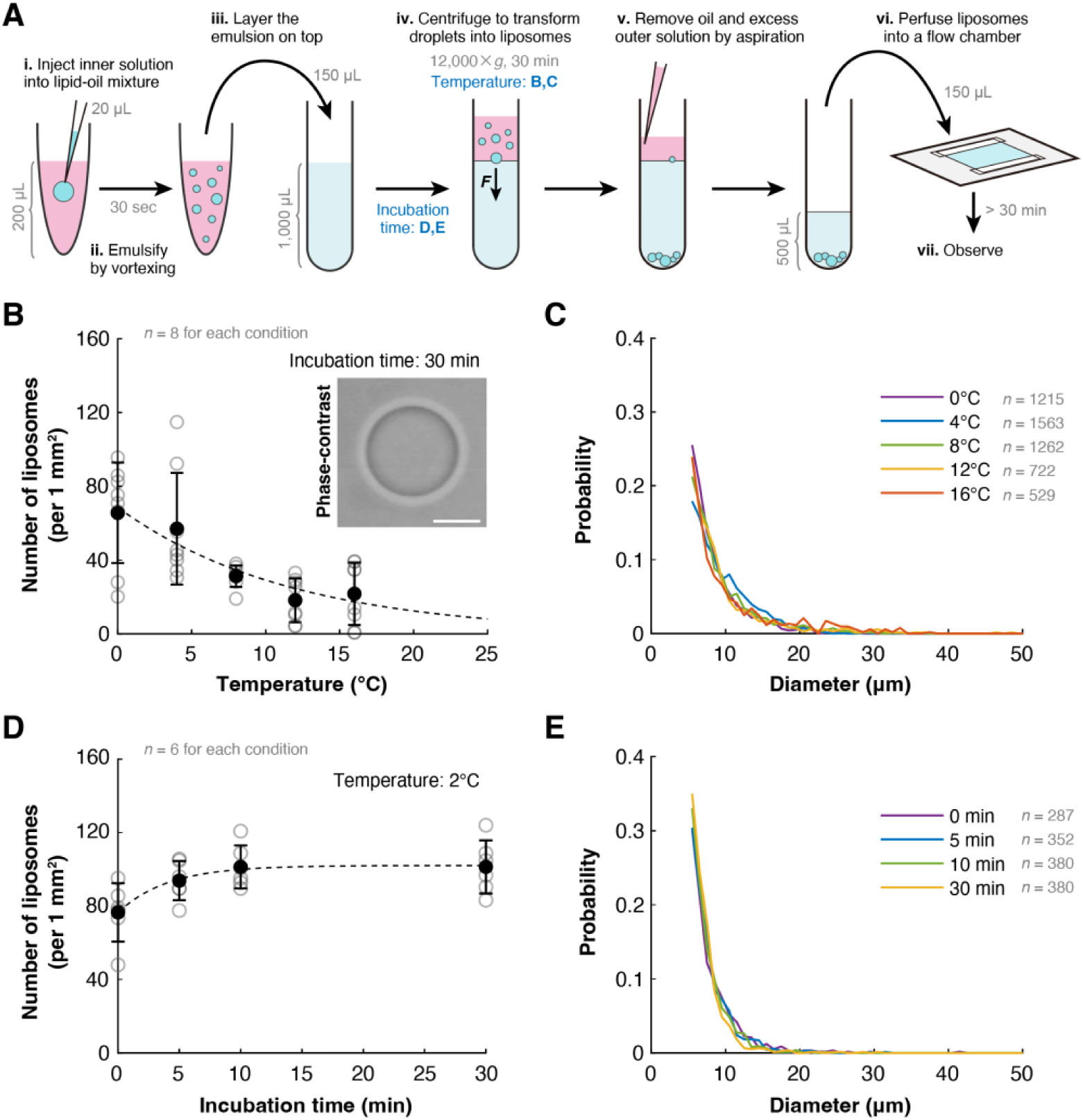
Effects of temperature and incubation time on the production yield and diameter of liposomes formed using the centrifugal sedimentation method. **A.** Schematic illustration of the centrifugal sedimentation method for liposome formation and observation. **B, C.** Effects of centrifugation temperature on (**B**) the production yield and (**C**) the diameter of liposomes. Liposomes larger than 5 μm in diameter were analyzed. (**B**) Closed circles and error bars indicate the means and standard deviations (SDs), respectively. Open circles represent the raw data. The broken line represents a fitting curve based on the equation *y* = *a* exp(−*bx*) with *a* = 68.6 and *b* = 0.0855. Eight independent experiments were performed for each condition. Inset: Representative phase-contrast image of a liposome (scale bar: 10 μm). **D, E.** Effects of incubation time before centrifugation on (**D**) the production yield and (**E**) the diameter of liposomes. Liposomes larger than 5 μm in diameter were analyzed. (**D**) Closed circles and open circles indicate the means and the raw data, respectively. The broken line represents a fitting curve based on the equation *y* = *c* − *a* exp(−*bx*) with *a* = 25.6, *b* = 0.249, and *c* = 102. Two independent experiments were performed for each condition.

**Figure 5.**
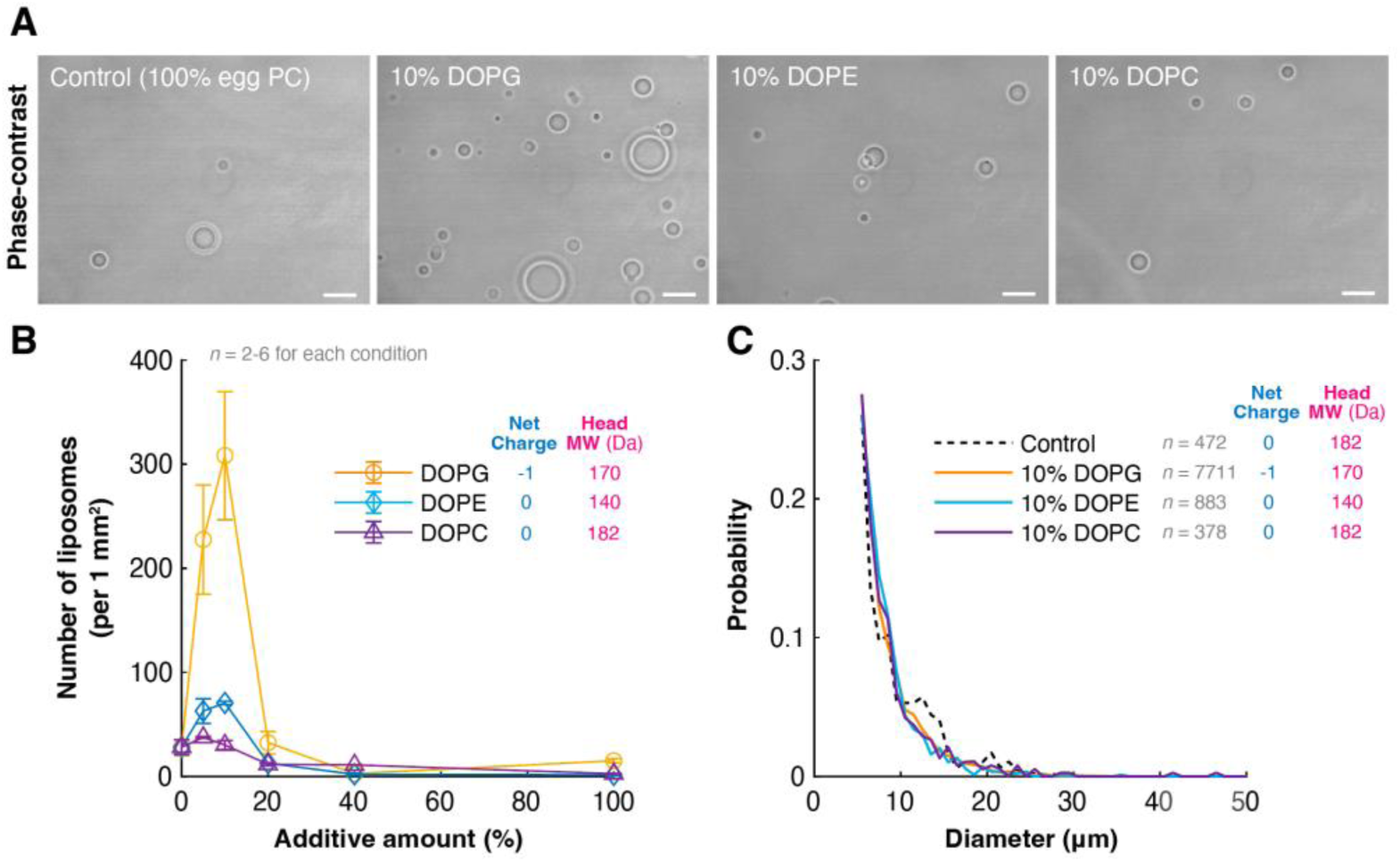
Effects of lipid composition on the production yield and diameter of liposomes formed using the centrifugal sedimentation method. **A.** Representative images of liposomes formed using the centrifugal sedimentation method. The liposomes were observed using a phase contrast microscope. Scale bars: 20 μm. **B, C.** Effects of lipid components and their additive amounts on (**B**) the production yield and (**C**) the diameter of liposomes. Liposomes larger than 5 μm in diameter were analyzed. Plots and error bars in (**B**) indicate the means and standard deviations (SDs), respectively. The incubation time before centrifugation (between step **iii.** and step **iv.** in **Fig. 4A**) and centrifugation temperature were fixed at 0 min and 4°C, respectively. Two to six independent experiments were performed for each condition. The net charges and molecular weights of the hydrophilic regions of the additive phospholipids or egg PC (control) are listed.

### Liposome formation using the natural sedimentation method

A 96-well microplate (*φ* = 6.7 mm well diameter; 92696, TPP) was used for liposome formation and observation using the natural sedimentation method (**Fig. 2A**). All procedures were performed at room temperature. First, 50 μL of the outer solution (A50 buffer containing 500 mM glucose) was added to each well. Next, 50 μL of the lipid-oil mixture was carefully layered on top (**Fig. 2A**, step i) and incubated for >10 min to assemble a lipid monolayer at the outer solution/oil interface. After the incubation, 50 μL of the lipid-oil mixture was transferred to a 1.5 mL plastic test tube (0030120086, Eppendorf), and then 5 μL of the inner solution (A50 buffer containing 150 mM sucrose, 350 mM glucose, and 1/1000 volume of 10 mM octadecyl rhodamine B chloride (R18) dissolved in DMSO) was injected to the lipid-oil mixture (**Fig. 2A**, step ii). The test tube was then vortexed for 30 sec using a vortex mixer (SI-0236, Scientific Industries) at maximum power to generate water-in-oil droplets (**Fig. 2A**, step iii). Immediately after the droplet formation, 20 μL of the emulsion was taken from the upper layer and then gently placed on top of the lipid monolayer formed in the microplate (**Fig. 2A**, step iv). The microplate was then incubated for >20 min to transform droplets into liposomes (**Fig. 2A**, step v). During incubation, the microplate was kept horizontal and protected from vibrations. After incubation, the microplate was carefully mounted on an inverted microscope and liposomes were observed (**Fig. 2A**, step vi).

### Liposome formation using the centrifugal sedimentation method

Liposomes were generated according to our previous report <u>(Chiba 2014)</u> (See **Fig. 4A**). First, 200 μL of the lipid-oil mixture was transferred to a 1.5 mL plastic test tube (0030120086, Eppendorf) and incubated on ice (**Fig. 4D,E**, **Fig. 5**, **Fig. 6**) or at various temperatures inside a refrigerated centrifuge (CF-15R, Hitachi) (**Fig. 4B,C**) for 15 min. After the incubation, 20 μL of the inner solution (A50 buffer containing 150 mM sucrose, 350 mM glucose), preincubated at the same temperature, was added (**Fig. 4A**, step i), then the test tube was vortexed for 30 sec using a vortex mixer (SI-0236, Scientific Industries) at maximum power to generate water-in-oil droplets (**Fig. 4A**, step ii). After the droplet formation, the test tube was incubated on ice (**Fig. 4D,E**, **Fig. 5**, **Fig. 6**) or at various temperatures inside a refrigerated centrifuge (CF-15R, Hitachi) (**Fig. 4B,C**) for 5 min. After the incubation, 150 μL of the emulsion was taken from the upper layer and then gently placed on top of 1 mL of the outer solution (A50 buffer containing 500 mM glucose) in a 1.5 mL glass test tube (0407-03, Maruemu), which was preincubated at the same temperature for >20 min (**Fig. 4A**, step iii). The test tube was then incubated on ice (**Fig. 4D,E**, **Fig. 5**, **Fig. 6**) or at various temperatures inside a refrigerated centrifuge (CF-15R, Hitachi) (**Fig. 4B,C**) for various duration times (**Fig. 4D,E**), 30 min (**Fig. 4B,C**), or 0 min (**Fig. 5**, **Fig. 6**) to assemble a lipid monolayer at the water/oil interface. After the incubation, the test tube was centrifuged at 12,000 ×*g* for 30 min at various temperatures (**Fig. 4B,C**), 2°C (**Fig. 4D,E**), or 4°C (**Fig.5**, **Fig.6**) (**Fig. 4A**, step iv). Meanwhile, an observation chamber was assembled by placing two double-sided tapes (thickness 0.3 mm; PBW-20, 3M) onto a silicone-coated coverslip (36 × 24 mm^2^; custom-ordered, Matsunami) with a coverslip (18 × 18 mm^2^; C218181, Matsunami) on top. The inner distance between the two tapes was ∼8 mm, and the inner volume of the chamber was ∼40 μL. First, the flow chamber was coated with 60 μL of Pluronic-F127 (10 mg/ml dissolved in A50 buffer) for >1 min to coat the surface. Then, the flow chamber was washed with 600 μL of the outer solution (A50 buffer containing 500 mM glucose). After the centrifugation, the oil layer and the upper half layer of the outer solution were removed by aspiration (**Fig. 4A**, step v). Then, the remaining liposome solution (∼500 μL) was mixed gently by pipetting 30 times to disperse liposomes homogeneously, and 150 μL was perfused into the flow chamber (**Fig. 4A**, step vi). The flow chamber was sealed with Valap and kept horizontal for >30 min at room temperature to settle down liposomes on the bottom coverslip. Finally, liposomes were observed under a microscope (**Fig. 4A, s**tep vii). Note that, in the experiments examining temperature dependence (**Fig. 4B,C**), the temperature of the centrifuge rotor was measured before and after the centrifugation using a K-type thermocouple and verified if the temperature was within ± 2°C of the set temperature in each experiment. Liposomes produced out of this temperature range were eliminated from the analysis.

**Figure 6.**
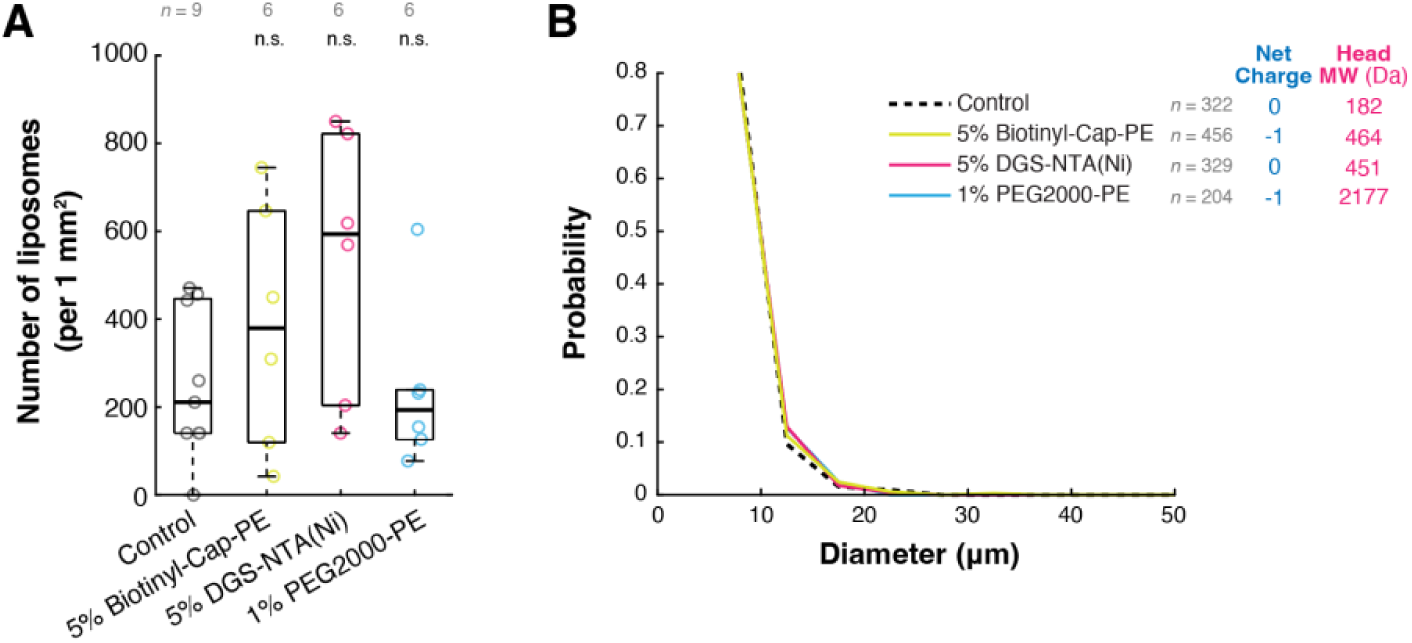
Effects of large head lipids on the production yield and diameter of liposomes formed using the centrifugal sedimentation method. **A.** Number of liposomes observed in 1 mm^2^ field of view. The raw data are overlaid on the box plot as a scatter plot. Liposomes larger than 5 μm in diameter were analyzed. The incubation time before centrifugation (between step **iii.** and step **iv.** in **Fig. 4A**) and centrifugation temperature were fixed at 0 min and 4°C, respectively. Six to nine independent experiments were performed for each condition. n.s.: *p* ≥ 0.05 (Welch’s *t*-test, two-sided). **B.** Distribution of the liposome diameters observed in (**A**). The net charges and molecular weights of the hydrophilic regions of the additive phospholipids or egg PC (control) are listed.

### Water-in-oil droplet formation for observation

First, 50 μL of the lipid-oil mixture was transferred to a 1.5 mL plastic test tube (0030120086, Eppendorf), and then of 5 μL of the inner solution (A50 buffer containing 150 mM sucrose, 350 mM glucose, and 1/1000 volume of 10 mM octadecyl rhodamine B chloride (R18) dissolved in DMSO) was added. The test tube was then vortexed for 30 sec using a vortex mixer (SI-0236, Scientific Industries) at maximum power to generate water-in-oil droplets. Immediately after the droplet formation, 20 μL of the emulsion was taken from the upper layer and then diluted 5 times with 80 μL of the lipid-oil-mixture. After the dilution, 20 μL of the droplet solution was perfused into a flow chamber, assembled by placing two double-sided tapes (thickness 0.1 mm; NW-20, Nichiban) onto a silicone-coated coverslip (36 × 24 mm^2^; custom-ordered, Matsunami) with a coverslip (18 × 18 mm^2^; C218181, Matsunami) on top. The flow chamber was sealed with Valap, and then droplets were observed under a microscope.

### Microscopy

Epi-fluorescence and bright-field images of liposomes prepared by the natural sedimentation method and water-in-oil droplets (**Figs. 2 and 3**) were observed by an inverted microscope (custom-built) equipped with ×40 objective (LCPlanFI 40×/0.60 Ph2, Olympus), an electron multiplying charge coupled device (EM-CCD) camera (iXon3, Andor Technology), and a motorized sample stage (Custom-built, OptoSigma). Phase-contrast images of liposomes prepared by the centrifugal sedimentation method (**Figs. 4 and 5**) were observed by an inverted microscope (DIAPHOT 300, Nikon) equipped with ×60 objective (PlanApo NA 1.40 oil Ph4), CCD camera (CCD-300-RCX, Dage-MTI), and a motorized sample stage (Custom-built, OptoSigma). Bright-field images of liposomes prepared by the centrifugal sedimentation method (**Fig. 6**) were observed by an inverted microscope (IX73, Olympus or custom-built) equipped with ×60 objective (PlanApo 60×/1.40, Olympus), an electron multiplying charge-coupled device (EM-CCD) camera (iXon3, Andor Technology), and a motorized sample stage (Custom-built, OptoSigma).

### Image analysis

Bright-field images of liposomes and droplets (**Figs. 2,3, and 6**) were scanned along the Z-axis from – 10 to 100 μm at 1-μm intervals (0 μm corresponds to the position of the bottom coverslip) across 5 different fields of view (one field of view: 172.9 × 172.9 μm^2^) (**Figs. 2 and 3**) or 10 different fields of view (one field of view: 119.4 × 119.4 μm^2^) (**Fig. 6**), at room temperature. Spherical liposomes with a diameter larger than 5 μm were manually detected, and their numbers and diameters were measured using Fiji/Image J (NIH). Alternatively, phase-contrast images of liposomes (**Figs. 4 and 5**) were scanned along the Z-axis from –25 to 100 μm at 0.67-μm intervals, across 40 (**Fig. 4D,E**) or 200 (**Fig. 4B,C**, **Fig. 5**) different fields of view (one field of view: 204.2 × 153.1 μm^2^), at room temperature. Spherical liposomes with a diameter larger than 5 μm were automatically detected, and their numbers and diameters were measured using a custom-built program written in LabVIEW (National Instruments) <u>(Chiba 2014)</u>.

### Statistical analysis

The two-sided Welch’s *t*-test was performed using MATLAB (MathWorks).

### Reproducibility

All experiments were repeated at least two times to confirm reproducibility. The same production lots of phospholipids and mineral oil were consistently used throughout each series of experiments. The variation in the number of liposomes under the control condition (100% egg PC) across different experimental series is likely attributable primarily to differences between the production lots.

## RESULTS

### Effects of lipid composition on the production yield of liposomes formed using the natural sedimentation method

We first examined the effects of lipid composition on the number of liposomes produced using the natural sedimentation method (**Fig. 2A**). All the following procedures were carried out at room temperature. Briefly, the outer solution was added to a microwell of a 96-well polystyrene microplate, and then 1 mM phospholipids dissolved in mineral oil were carefully layered on top (**Fig. 2A**, step i) and incubated for more than 10 minutes to allow the assembly of a lipid monolayer at the water/oil interface. After incubation, the inner solution containing 10 μM R18 membrane staining dye was injected into the lipid-oil mixture in a test tube (**Fig. 2A**, step ii), and the tube was vortexed for 30 seconds to generate water-in-oil droplets (**Fig. 2A**, step iii). Immediately after droplet formation, the emulsion was gently placed on top of the lipid monolayer formed in the microplate (**Fig. 2A**, step iv). The microplate was then incubated for more than 20 minutes to allow the spontaneous transformation of droplets into liposomes due to the mass density difference between the inner solution and mineral oil, followed by sedimentation of liposomes to the bottom surface due to the mass density difference between the inner and outer solutions (**Fig. 2A**, step v). The mass density difference between the inner solution and the lipid-oil mixture and between the inner and outer solutions was estimated to be 282 mg/mL and 24.3 mg/mL, respectively, based on the molecular weights of the chemical components. During incubation, the microplate was kept horizontal and protected from vibrations. After incubation, the microplate was carefully mounted on an inverted microscope, and liposomes were observed (**Fig. 2A**, step vi). We located the center of the microwell and focused on the bottom surface using the epi-fluorescence images (**Fig. 2B**), and then defined the origin of the XYZ-axis of the motorized stages. After that, we scanned epi-fluorescence (**Fig. 2B**) and bright-field (**Fig. 2C**) images of liposomes along the Z-axis from –10 μm to 100 μm at 1-μm intervals, both at the center of the microwell (X, Y) = (0 μm, 0 μm) and at four positions located 0.84 mm from the center, specifically at (X, Y) = (±600 μm, ±600 μm). Then, we manually detected liposomes using bright-field images with the aid of fluorescence images, and counted the numbers and measured the diameters of spherical liposomes larger than 5 μm.

First, we compared the number of liposomes formed using 100% egg PC (control) and a 90:10 molar ratio mixture of egg PC and DOPC. We confirmed that replacing 10% of egg PC with DOPC did not significantly affect the number of liposomes (**Fig. 2D**, gray and purple plots). In other words, a 10% replacement of a mixture of unsaturated and saturated fatty acids (**Fig. S1**) with 18:1-18:1 unsaturated fatty acid does not influence the production yield of liposomes. Next, we replaced 10% of egg PC with 18:1-18:1 unsaturated fatty acid lipids containing different head groups (DOPE, DOPA, DOPG, and DOPS) and examined the effects. We observed that replacement with DOPE led to a slight increase in both the mean and median number of liposomes (**Fig. 2D**, blue plots), but the difference was not statistically significant (*p* = 0.06; Welch’s *t*-test, two-sided). By contrast, replacement with DOPA, DOPG, or DOPS drastically increased the number of liposomes by 13.7-, 8.83-, and 8.84-fold, respectively, as mean values compared to the control (100% egg PC) (**Fig. 2D**, green, yellow, and orange plots). These three lipids share a common property: each carries a single negative charge on its hydrophilic head region. This suggests that charged lipid head groups enhance the production yield of liposomes. In addition to electric charge, the molecular weight of the hydrophilic head region is also an essential factor that characterizes the physical properties of the membrane <u>(Boal 2012, Phillips 2013)</u>. Interestingly, while the head regions of PG and PS have a similar molecular weight to PC, the molecular weight of PA’s head region is nearly half that of PC, and the number of liposomes seems to be the highest among the three charged head groups. Specifically, DOPA increased the number of liposomes by 1.56-fold and 1.55-fold compared to DOPG and DOPS, respectively, as mean values, and *p* = 0.06 between DOPA and DOPG, *p* = 0.02 between DOPA and DOPS (Welch’s *t*-test, two-sided). Moreover, PE, whose head group has a molecular weight 77% smaller than that of PC, showed a similar trend, with a slight increase in liposome number. These observations suggest that, in addition to electric charge, a smaller head group also contributes to enhancing the liposome production yield.

We next investigated whether a larger head group enhances the production yield of liposomes. To this end, we examined the effects of functionalized lipids with 18:1-18:1 unsaturated fatty acid, which have been used for protein localization on membranes, namely phospholipids conjugated with biotin (Biotinyl-Cap-PE) or Ni-NTA (DGS-NTA(Ni)) <u>(Carvalho 2013, Peters 2015, Loiseau 2016, Dürre 2018, Buddingh 2020, Razavi 2024, Sakamoto 2024, Monck 2024, Sakamoto 2024)</u>, as well as lipids used for membrane passivation, namely phospholipids conjugated with PEG (PEG2000-PE) <u>(Carvalho 2013, Loiseau 2016, Dürre 2018, Razavi 2024, Sakamoto 2024)</u>. The molecular weight of the hydrophilic head region is nearly double (Biotinyl-Cap-PE and DGS-NTA(Ni)) or more than 10 times larger (PEG2000-PE) compared to PC. We replaced egg PC with these functionalized lipids at typical percentages used in biophysical and synthetic biology experiments. Replacing 5% (for Biotinyl-Cap-PE and DGS-NTA(Ni)) or 1% (for PEG2000-PE) of egg PC with these functionalized lipids significantly increased the number of liposomes, by 3.21-, 3.21-, and 2.96-fold, respectively, as mean values compared to the control (100% egg PC) (**Fig. 2D**, lime, magenta, and cyan plots). This result suggests that a larger head group also facilitates liposome production yield. Note that, while Biotinyl-Cap-PE and PEG2000-PE carry a negative electric charge, DGS-NTA(Ni) has no net electric charge. However, the effect of electric charge was undetectable under the current conditions (*p* = 1.0 between DGS-NTA(Ni) and Biotinyl-Cap-PE, *p* = 0.67 between DGS-NTA(Ni) and PEG2000-PE, two-sided Welch’s *t*-test). Therefore, in the case of functionalized lipids, the large head size predominantly contributes to enhancing liposome production yield.

Finally, we examined the effects of a lipid conjugated with a fluorescent dye. Among the various commercially available fluorescent lipids, we selected ATTO488-conjugated lipid (ATTO488-PE) as a representative <u>(Jahnke 2022, Jahnke 2022, Drabik 2024)</u>. According to the statistical test, we did not detect a significant increase in the number of liposomes when 1% of egg PC was replaced with ATTO488-PE (**Fig. 2D**, brown plots), although the mean number of liposomes increased by 1.37-fold compared to the control (100% egg PC). This is presumably due to the lower replacement percentage (1%) compared to lipids with similar head sizes, such as Biotinyl-Cap-PE and DGS-NTA(Ni), which were used at 5% (**Fig. 2D**, lime and magenta plots). Collectively, we systematically investigated the effects of lipid composition on the number of liposomes produced using the natural sedimentation method, and found that the addition of charged head lipids, as well as larger or smaller head lipids, enhances the liposome production yield.

### Effects of lipid composition on the size of liposomes formed using the natural sedimentation method

We next investigated whether lipid composition modulates the size of liposomes produced by the natural sedimentation method. We quantified the probability distributions of liposome diameters under each condition in **Fig. 2D** and compared them with the control distribution (100% egg PC). We found that, when comparing lipids with similar or smaller head sizes, the electronic charge of the head region— whether neutral or negative—had no apparent effect on liposome size (**Fig. 3A**, left). In contrast, lipids with larger head groups tended to shift the liposome size distribution toward larger sizes, regardless of their electronic charge (**Fig. 3A**, right). This result suggests that replacing a small fraction of PC with a larger head lipid can increase liposome size. To further characterize this effect, we calculated the probability of liposomes larger than 10 μm in diameter under each condition and plotted these values against the molecular weight of the additive lipid head (**Fig. 3B**, left). We found a strong positive correlation between the molecular weight of the additive lipid head and the probability of liposomes larger than 10 μm in diameter (**Fig. 3B**, left). Furthermore, we categorized these liposomes into three size groups: 10–20 μm, 20–30 μm, and larger than 30 μm, and compared their distributions with the molecular weight of the additive lipid head (**Fig. 3B**, right). As a result, we observed positive correlations in all three groups, indicating that lipids with larger head groups can produce larger liposomes.

To dissect the mechanism underlying the observed phenomenon, we examined whether template droplets increased in size before transforming into liposomes or whether larger droplets exhibited enhanced transformation efficiency, in the presence of larger head lipids. To this end, we observed water-in-oil droplets using a bright-field microscope immediately after the formation (**Fig. 3C**) and measured their size distribution, as we did for liposomes. The analysis revealed that the size distributions were comparable across conditions, including smaller head lipids (10% DOPE and 10% DOPA), larger head lipids (5% DGS-NTA(Ni) and 1% PEG2000-PE), and the control (100% egg PC) (**Fig. 3D**). This result indicates that replacing PC with different head size lipids does not alter droplet size within this additive amount range (maximum 10%) and diameter range (larger than 5 μm). This finding suggests that large head lipids enhance the transformation efficiency of water-in-oil droplets into liposomes, rather than altering droplet size.

To gain further insights into the mechanism, we estimated the droplet-to-liposome transformation efficiency across various droplet diameters and compared the values among different lipid compositions. Specifically, we divided the probability distribution of liposome diameter **(Fig. 3A**) by that of droplet diameter (**Fig. 3D**) and then normalized the resulting probabilities against the diameter. Interestingly, while the droplet-to-liposome transformation efficiency exhibited a single peak at the 15– 20 μm or 20–25 μm diameter bin in the control condition (100% egg PC) and with 10% DOPE, 10% DOPA, and 5% DGS-NTA(Ni), the relative transformation efficiency showed the highest peak at the 30–35 μm diameter bin in the presence of 1% PEG2000-PE (**Fig. 3E**). This result further supports the idea that larger head lipids promote the formation of larger liposomes by enhancing the transformation efficiency of larger water-in-oil droplets.

### Effects of temperature and incubation time on the production yield and diameter of liposomes formed using the centrifugal sedimentation method

Although the natural sedimentation method has been widely used <u>(Takiguchi 2008, Yanagisawa 2011, Takiguchi 2011, Ito 2013)</u>, the centrifugal sedimentation method is more commonly employed in biophysical and synthetic biology experiments. In these experiments, a mixture of purified proteins <u>(Tsai 2011, Maeda 2012, Carvalho 2013, Peters 2015, Loiseau 2016, Nies 2018, Dürre 2018, Berhanu 2019, Bashirzadeh 2021, Buddingh 2020, Razavi 2024, Sakamoto 2024, Matsubayashi 2024, Sakamoto 2024)</u> or cell extracts <u>(Noireaux 2004, Takamori 2025)</u>, as well as a mixture of protein expression system and protein-encoded mRNAs <u>(Noireaux 2004, Berhanu 2019)</u> are encapsulated into water-in-oil droplets and the droplets are transformed into liposomes using the centrifugal sedimentation method. After liposome formation, the liposomes are transferred to an observation chamber. Then, the sample temperature is elevated to activate enzymatic reactions and observe reaction dynamics and morphological changes under an optical microscope. In such experiments, it is generally essential to maintain a lower temperature during liposome production to suppress enzymatic activity and minimize the duration of these procedures for practical applications. Otherwise, the reactions of interest may be completed before observation can begin. To date, the effects of temperature and incubation time on the lipid monolayer assembly process at the oil/outer solution interface have been characterized using synthetic POPC as a representative lipid <u>(Moga 2019)</u>. However, little is known about how these parameters influence the production yield and diameter of egg PC-based liposomes, a major lipid used for biophysical and synthetic biology experiments <u>(Takiguchi 2008, Yanagisawa 2011, Takiguchi 2011, Loiseau 2016, Dürre 2018, Razavi 2024, Sakamoto 2024, Sakamoto 2024)</u>. Since egg PC is a mixture of various unsaturated and saturated phosphatidylcholine lipids (**Fig. S1**), it is expected to retain the complex temperature-dependent physical properties of biological membranes <u>(Marsh 2013)</u>.

Here, we examined the effects of temperature on the production yield and diameter of liposomes using 100% egg PC as a representative lipid component. We prepared water-in-oil droplets at various temperatures (**Fig. 4A**, steps i and ii), layered the emulsion onto the outer solution preincubated at the same temperature (**Fig. 4A**, step iii), and then centrifuged it at 12,000 × g for 30 min to transform the droplets into liposomes at various temperatures (**Fig. 4A**, step iv). After the centrifugation, we removed the oil layer and half the volume of the outer solution by aspiration. Then, we dispersed liposomes sedimented at the bottom of the glass test tube by gentle pipetting, and the liposome solution was perfused into a flow chamber. The liposomes were observed under a phase-contrast microscope (**Fig. 2B**, inset). Spherical liposomes larger than 5 μm in diameter were automatically detected using a custom-built software, and the numbers were counted while the diameters were measured (**Fig. S2**). We found that the number of liposomes monotonically increased as the temperature decreased from 16°C to 0°C (**Fig. 4B**). This observation is contrary to a previous report using POPC, in which the yield appears to increase marginally with temperature <u>(Moga 2019)</u>. This discrepancy might reflect the complex temperature-dependent nature of egg PC (**Fig. S1**). We also found that the size distributions are comparable between the temperatures within this range (**Fig. 4C**).

We next investigated the effect of incubation time on the lipid monolayer assembly process at the oil/water interface. In this experiment, the centrifugation process (**Fig. 4A**, step iv) was carried at 2°C and all other liposome production procedures (**Fig. 4A**, steps i-iii) were carried out on ice. After generating water-in-oil droplets (**Fig. 4A**, steps i and ii), the emulsion was layered onto the outer solution preincubated on ice (**Fig. 4A**, step iii). The sample was then incubated for various durations before centrifugation (**Fig. 4A**, step iv). After centrifugation, the liposomes were collected and observed under a microscope (**Fig. 4A**, steps v, vi, and vii). We found that the number of liposomes slightly increased as the incubation time increased from 0 to 10 minutes and then reached a plateau (**Fig. 4D**), indicating that a 10-minute incubation is sufficient for lipid monolayer assembly. We also found that the size distribution was not affected by incubation time (**Fig. 4E**). Overall, our observations revealed that a lower temperature is optimal for the production of egg PC-based liposomes, and a 10-minute incubation is sufficient for lipid monolayer assembly at the oil/water interface on ice.

### Effects of lipid composition on the production yield and diameter of liposomes formed using the centrifugal sedimentation method

We next investigated the effects of lipid composition on the number and diameter of liposomes using the centrifugal sedimentation method (**Fig. 4A**). In the following experiments, the centrifuge temperature and incubation time were fixed at 4°C and 30 minutes, respectively, and all other liposome production procedures were carried out on ice. As observed in the natural sedimentation method (**Fig. 2D** and **Fig. 3A**), replacing PC with charged lipids or smaller head lipids was expected to enhance production yield without altering the liposome size distribution. In this experiment, we selected DOPG as a representative charged lipid with a similar head size and DOPE as a representative non-charged lipid with a smaller head size, then varied their additive percentages from 2.5% to 100%.

We found that lipid replacement enhanced the number of liposomes, reaching a peak at 10% replacement for both DOPG and DOPE, yet further replacement drastically decreased the number of liposomes (**Fig. 5B**, orange and blue plots). Replacing egg PC with DOPC did not increase the number of liposomes at any tested percentage (**Fig. 5B**, purple plots), confirming that the enhancement in production yield by DOPG and DOPE is attributable to their lipid head properties. We also confirmed that lipid replacement did not alter the liposome diameter distribution (**Fig. 5C**), which is consistent with the results of the natural sedimentation method (**Fig. 3A**).

### Effects of large head lipids on the production yield and diameter of liposomes formed using the centrifugal sedimentation method

Finally, we examined whether large head lipids promote the production yield and formation of larger liposomes by the centrifugal sedimentation method, as observed in the natural sedimentation method (**Fig.2**, **Fig. 3**). Specifically, we quantified the number and diameter of liposomes produced using three different large head lipids (Biotinyl-Cap-PE, DGS-NTA, and PEG2000-PE) and compared the results with the control (**Fig. 6**). In contrast to the natural sedimentation method, no statistically significant difference in production yield was observed between the control and any of the three conditions (**Fig. 6A**). Similarly, no significant difference in liposome diameter distributions were observed among the conditions (**Fig. 6B**). Taken together, our results indicate that the presence of 1-5% large head lipids did not significantly increase either the production yield or the diameter of liposomes in the centrifugal sedimentation method.

## DISCUSSION

In synthetic biology research, phosphatidylcholine (PC) is commonly used as the basal lipid, either as natural PC purified from chicken eggs or soybeans, or as synthetic lipids such as DOPC and POPC. These lipids are often mixed with negatively charged lipids, including DOPG, POPG, DOPS, and POPS, to reduce non-specific interactions between protein and the membrane, as typical proteins carry a net negative charge under physiological pH. Furthermore, it is widely accepted that the addition of a few to several tens of percent of charged lipids empirically enhances the production yield of liposomes. However, to the best of our knowledge, no direct evidence has been reported to support this assumption.

In this study, we systematically investigated the effects of lipid composition on the production yield of cell-sized liposomes and found that the addition of charged lipids promotes liposome formation using the inverted emulsion technique, both in the natural sedimentation method (**Fig. 2**, **Fig. 3**) and the centrifugal sedimentation method (**Fig. 4**, **Fig. 5**, **Fig. 6**). Using natural PC purified from chicken eggs (egg PC) as the basal component, we demonstrated that the addition of 10% DOPG, 10% DOPS, or 10% DOPA increased the production yield by ∼10-fold under the natural sedimentation method (**Fig. 2D**). This trend was also observed in the centrifugal sedimentation method: the addition of DOPG enhanced the production yield, with a maximum effect observed at 10% DOPG (∼10-fold increase; **Fig. 5B**). In addition to electric charge, we found that the molecular weight of the hydrophilic head group also influences liposome production yield. In the natural sedimentation method, the addition of lipids with either smaller or larger head groups marginally increased the number of liposomes (**Fig. 2D**). In the centrifugal sedimentation method, a similar trend was observed for a smaller head group, whereas no significant increase in the production yield was observed for larger head groups (**Fig. 5B**, **Fig. 6A**).

We also examined the effect of additive phospholipids on the size distribution of liposomes. Systematic analysis on the natural sedimentation method (**Fig. 3**) revealed that the formation probability of larger liposomes positively correlates with the molecular weight of the head group of the additive phospholipids (**Fig. 3A,B**). We confirmed that the diameter distribution of water-in-oil droplets does not differ significantly with the inclusion of lipids bearing larger or smaller head groups (**Fig. 3C,D**). These findings suggest that, although the underlying physical mechanism remains unresolved, the head group size of phospholipids modulates the optimal droplet diameter for efficient droplet-to-liposome transformation in the natural sedimentation method **(Fig. 3E**). However, in the centrifugal sedimentation method, we observed no significant difference in the size of liposomes produced using larger head groups (**Fig. 6B**), and their sizes were consistently smaller than those produced by natural sedimentation method including the control condition (**Fig. 3A,B**, **Fig. 5C**, **Fig. 6B**). These results suggest that the centrifugal force facilitates the passage of smaller droplets that may otherwise fail to cross the oil-water interface in the natural sedimentation method, thereby the effect of larger head groups may be masked by the strong centrifugal force.

Using egg PC, we also examined the effects of incubation time for lipid monolayer formation at the oil/outer solution interface and the centrifugation temperature. Pautot *et al*. previously reported <u>(Pautot 2003)</u> that lipid adsorption at the oil/water interface is a diffusion-limited process and appears to be more complex than that of typical surfactants. The authors also noted that achieving full monolayer coverage requires ∼30 minutes for charged lipids and ∼90 minutes for zwitterionic lipids. However, our results demonstrate that, at least for egg PC, a 10-minute incubation is sufficient for practical applications. Even without incubation, the production yield retains ∼75% of the maximum saturated value (**Fig. 4D**), and the size distribution of liposomes shows no significant differences across the tested incubation times (0-30 min) (**Fig. 4E**). Regarding temperature dependence, a previous study using POPC as the model phospholipid reported that increasing the temperature from 4 °C to 37 °C marginally enhanced the production yield by ∼1.4-fold, while the mean size of liposomes produced at 37 °C was slightly smaller than those produced at lower temperatures <u>(Moga 2019)</u>. In contrast, our results with egg PC show that lower temperatures enhance the production yield, with a ∼2-fold increase observed when the temperature is reduced from 16 °C to 0 °C (**Fig. 4B**). However, temperature did not have a detectable impact on the size distribution of liposomes within this range (**Fig. 4C**). This difference might primarily arise from the distinct temperature-dependent physical properties of POPC, a single-component lipid, versus egg PC, a heterogeneous mixture of PC species. The phase transition temperature (*T*_m_) from the ordered (gel) to disordered (liquid crystalline) phase for POPC is –9 °C <u>(Marsh 2013)</u>. In contrast, egg PC comprises a mixture of lipids with a broad range of *T*_m_ values <u>(Marsh 2013)</u>. Notably, at least nine potential lipid species present in egg PC exhibit *T*_m_ values that fall within the temperature range tested in our study (0-25 °C; **Fig. 4B,C**) <u>(Marsh 2013)</u>. These facts also imply that the optimal temperature for enhancing liposome production yield may be influenced by the physical properties of the additive lipids, particularly the acyl chain length and degree of unsaturation in their hydrophobic tails. Further studies are needed to elucidate how these structural features of phospholipids affect both the production yield and size distribution of liposomes generated by the inverted emulsion technique.

Based on the observations in the present study and in our previous report <u>(Chiba 2014)</u>, we propose three potential key factors to enhance the liposome production yield for the inverted emulsion method (**Fig. 7**). The first key factor is the abundant supply of lipid molecules (**Fig.7**, i). We previously reported that increasing the lipid concentration dissolved in mineral oil up to 5 mM increased the number of liposomes produced by the centrifugal sedimentation method <u>(Chiba 2014)</u>. This finding appears consistent with the observation in the present study that longer incubation times for lipid monolayer formation enhance the production yield. It is noteworthy that a further increase in lipid concentration did not enhance liposome production, but instead resulted in contamination with multilamellar liposomes <u>(Chiba 2014)</u>.

**Figure 7.**
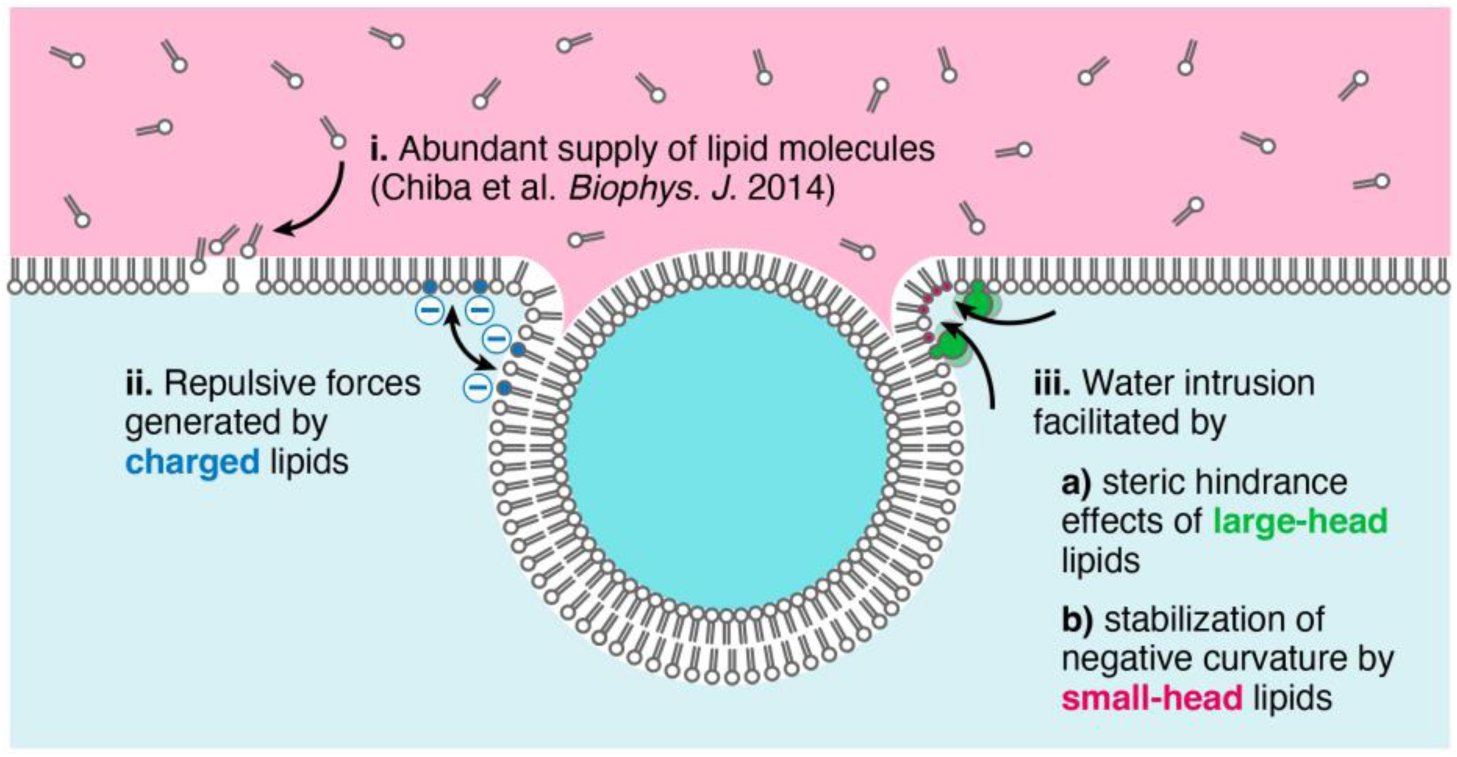
Potential key factors for enhancing liposome production yield. Three key factors may contribute to increasing the number of liposomes produced by the inverted emulsion method: **i.** The abundant supply of lipid molecules to the oil/water interface, demonstrated in our previous report <u>(Chiba 2014)</u>, **ii.** Repulsive forces generated by charged lipids between lipid membranes, and **iii.** Intrusion of water molecules into the “neck” region of a suspended liposome beneath the oil/water interface, facilitated by **a)** steric hindrance effects of large-head lipids or **b)** stabilization of strong negative curvature by small-head lipids. All three factors may contribute to reducing the width of the “neck” region, thereby facilitating the detachment of suspended liposomes from the oil layer.

The second key factor will be repulsive forces generated by charged lipids between a lipid bilayer on a liposome and a lipid monolayer at the interface between the outer solution and oil (**Fig. 7**, ii). In the present study, we have shown that the addition of charged lipids with the molecular weight of the hydrophilic region comparable to PC (DOPG and DOPS) significantly increases the production yield, both in the natural sedimentation method (**Fig. 2D**) and the centrifugal sedimentation method (**Fig. 5B**). The addition of charged lipids with a small-head group (DOPA) drastically enhances the production yield in the natural sedimentation method (**Fig. 2D**). The addition of charged lipids with large-head groups also slightly increases the production yield (Biotinyl-Cap-PE and PEG2000-PE) in the natural sedimentation method. By contrast, in the centrifugal sedimentation method, no significant increase of the production yield was observed with the large-head groups (**Fig. 6A**). These results indicate that the repulsive force between charged lipids with small-head groups or those of comparable size to PC facilitates the passage of water-in-oil droplets through the oil-water interface in both methods. However, this effect diminishes when using charged lipids with larger head groups in the centrifugal sedimentation method. Note that the repulsive forces may also act between the lipids aligned at the oil-water interface and the lipids dispersed in the oil phase. At optimal percentages of charged lipids, such repulsive interactions in the oil phase might help form the monolayer of lipids at the oil-water interface, thereby further enhancing the production yield.

The third key factor will be the intrusion of water molecules at the neck of the suspended liposomes at the oil/water interface, facilitated by a) large-head lipids or b) stabilization of negative curvature by small-head lipids (**Fig. 7**, iii). We have shown in the present study that the addition of large-head lipids with neutral electric charge (DGS-NTA(Ni)) increases the production yield in the natural sedimentation method (**Fig. 2D**). In contrast, no significant difference was observed in centrifugal sedimentation method (**Fig. 6A**), indicating that the influence of the head group on the production yields is smaller than that of the centrifugal force. The addition of slightly smaller head lipids with neutral electric charge (DOPE) seems to marginally increase the production yield, also it shows no statistical significance in the natural sedimentation method (*p* = 0.06) (**Fig. 2D**). This effect was enhanced in the centrifugal sedimentation method (**Fig. 5B**).

In summary, we systematically investigated the effects of lipid composition on the number and size of liposomes produced using two standard liposome formation methods based on the inverted emulsion technique, employing natural PC purified from chicken eggs, one of the most widely used phospholipids in artificial cell research, as the base lipid. Our quantitative analysis provides direct evidence that the addition of charged lipids and/or lipids with smaller head groups to egg PC enhances the production yield of liposomes under both the natural sedimentation method and centrifugal sedimentation method. Furthermore, we found that the incorporation of lipids with larger head groups promotes the formation of liposomes with greater diameters under the natural sedimentation method, but this effect is diminished in the centrifugal sedimentation method. These findings offer valuable guidelines for optimizing preparation protocols for artificial cell systems using the inverted emulsion technique, and may support the advancement of more sophisticated artificial cell models in future studies.

## DATA AND CODE AVAILABILITY

The raw data used for generating the plots are summarized in **Table S1**. Raw images are available from the corresponding author upon reasonable request.

## ACKNOWLEDGMENTS

The authors thank all laboratory members for their assistance with the experiments and discussions. This work was supported by JST PRESTO, Japan (Grant No. JPMJPR20ED to M.M.); Grant-in-Aid for Transformative Research Areas (A) (Grant No. 22H05171 to M.M.) from the Ministry of Education, Culture, Sports, Science, and Technology, Japan; The Hakubi Project of Kyoto University (to M.M.); and NINS Astrobiology Center program research (Grant No. AB0714 to M.M.).

## AUTHOR CONTRIBUTIONS

M.M. designed the research. H.S., H.M., K.G., and M.M. performed experiments, analyzed the data, and discussed the results. H.M. and M.M. wrote the manuscript.

---

**Figure S1.**
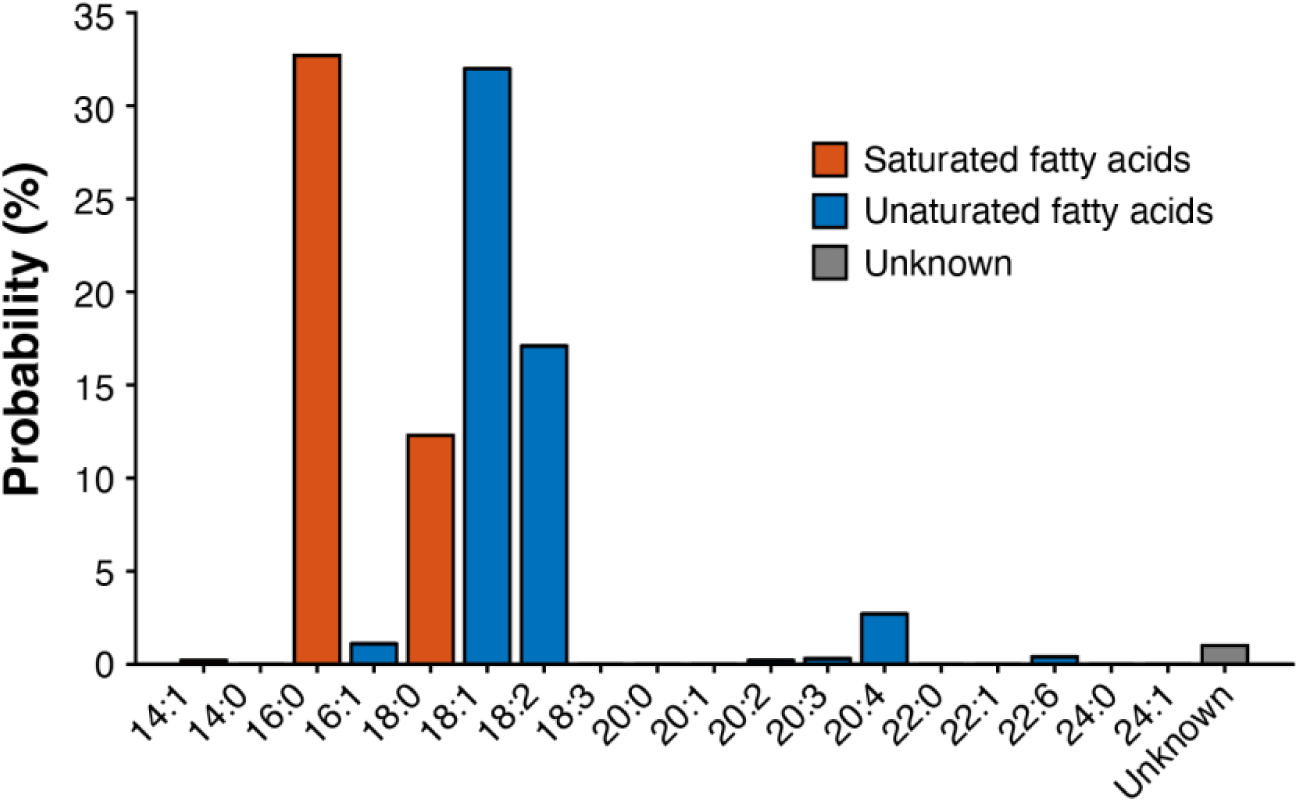
Fatty acid distribution of egg PC. The analysis data were downloaded from the webpage of Avanti Polar Lipids, Inc. Based on the fatty acid distribution, the mean molecular weight of the lipids was estimated to be 770.123 Da.

